# Attention induced perceptual traveling waves in binocular rivalry

**DOI:** 10.1101/2025.04.18.649496

**Authors:** João V.X. Cardoso, Hsin-Hung Li, David J. Heeger, Laura Dugué

**Affiliations:** Université Paris Cité, CNRS, Integrative Neuroscience and Cognition Center, F-75006 Paris, France; Department of Psychology, The Ohio State University, Columbus, OH 43201, USA; Department of Psychology, New York University, New York, NY 10003, United States; Center for Neural Science, New York University, New York, NY 10003, United States; Institut Universitaire de France (IUF), Paris, France

**Author notes:** **Corresponding author** João V.X. Cardoso.

**Keywords:** Attention, Binocular Rivalry, Computational Model, Traveling Waves

## Abstract

Cortical traveling waves –smooth changes of phase over time across the cortical surface– have been proposed to modulate perception periodically as they travel through retinotopic cortex. Yet, little is known about the underlying computational principles. Here, we make use of binocular rivalry, a perceptual phenomenon in which perceptual (illusory) waves are perceived when a shift in dominance occurs between two rival images. First, we assessed these perceptual waves using psychophysics. Participants viewed a stimulus restricted to an annulus around fixation, with orthogonal orientations presented to each eye. The stimulus presented to one eye was of higher contrast thus generating perceptual dominance. When a patch of higher contrast was flashed briefly at one position in the other eye, it created a change in dominance that started at that location of the flash and expanded progressively, like a wave, as the previously suppressed stimulus became dominant. We found that the duration of the perceptual propagation increased with both distance traveled and eccentricity of the annulus. Diverting attention away from the annulus reduced drastically the occurrence and the speed of the wave. Second, we developed a computational model of traveling waves in which competition between the neural representations of the two stimuli is driven by both attentional modulation and mutual inhibition. We found that the model captured the key features of wave propagation dynamics. Together, these findings provide new insights into the functional relevance of cortical traveling waves and offer a framework for further experimental investigation into their role in perception.

## Introduction

Cortical traveling waves have been observed across cortical areas associated with various sensory, motor, and cognitive processes and in various animal species (for review, Muller et al., 2018), suggesting a fundamental principle of brain function. In humans, waves of activity have been mostly described at a global scale, i.e., across a large portion of cortex (Hangya et al., 2011; Hindriks et al., 2014), using various invasive (electro-corticography, ECoG, Muller et al., 2016, and stereotactic electro-encephalography, sEEG, Alexander et Dugué, 2024) and non-invasive neuroimaging (electro-encephalography, EEG, Hughes, 1995, Alamia & VanRullen, 2019, and magneto-encephalography, MEG, Alexander et al., 2013) techniques. Mesoscopic waves, traveling across single brain regions, however, are far less investigated in humans as they require high spatial resolution beyond that of current neuroimaging tools (Petras et al., 2025; Grabot et al., 2024). Here, we aimed to fill this gap using psychophysics in human, combined with computational modeling. Specifically, we assessed the mechanistic role of mesoscopic traveling waves in perception.

Two prior psychophysics studies have assessed mesoscopic traveling waves non-invasively in human (Sokoliuk & VanRullen, 2016; Fakche & Dugué, 2024). They used a disk (inducer) oscillating in luminance at low frequencies (between 4 and 10Hz) and presented in the periphery of the visual field to induce brain oscillations at the corresponding frequencies and retinotopic cortical location. They showed that the perception of a low-contrast stimulus presented at nearby locations was modulated periodically at the inducer frequency, creating *induced perceptual cycles*. Critically, the phase of the inducer leading to the highest target detection performance shifted as a function of distance between the inducer and the target. These results support the idea that activity propagating within visual cortex, leads to corresponding perceptual consequences, i.e., the propagation of perceptual cycles across retinotopic visual space. In these studies, however, the spatio-temporal characteristics of the propagating activity within retinotopic cortex were determined by the oscillating disc, leaving unknown the spontaneous dynamic properties of brain activity. Additionally, the computational mechanisms underlying the propagation of cortical activity at the mesoscopic scale for perception remain unknown.

Here, we make use of a perceptual phenomenon in binocular rivalry (Levelt, 1965; Logothetis, 1998; Brascamp et al. 2005; Pearson & Brascamp, 2008; Blake & O’Shea, 2009; Blake, 2022), in which, under certain conditions, perceptual (illusory) waves are perceived when a shift in dominance occurs between two rival images (Wilson et al., 2001). Previous functional Magnetic Resonance Imaging (fMRI) studies have shown that such illusory perceptual changes, although present under constant physical stimulation, are correlated with the propagation of cortical activity in the primary visual cortex (V1; Wilson et al., 2001; Lee et al., 2003; Lee et al., 2005; Lee et al., 2007).

Building on these findings, we proposed novel psychophysical experiments designed to measure previously uncharacterized properties of perceptual waves, such as intra-individual variations in propagation speed across different stimulus eccentricities, ranging from foveal to peripherical areas. We subsequently developed a computational model of binocular rivalry that captures the expected cortical dynamics of these waves, enabling the investigation of how attentional modulation and eccentricity influence their propagation.

The model is an extension of a previously proposed model of binocular rivalry (Li et al., 2017). It relied on covert, exogenous (involuntary) attention modeled as multiplicative gains, and was able to explain several perceptual phenomena associated with binocular rivalry, such as its attention-dependent dynamics (Brascamp & Blake, 2012; Zhang et al., 2011). Here we introduce a spatial structure to examine how attentional modulation influences the dynamics of traveling waves within the model. The modified model posits that attention amplifies the neural representation of the dominant percept while attenuating that of the suppressed percept, thus prolonging the duration of the dominant percept and decelerating wave propagation. We calibrated the model against psychophysics data measuring the speed of traveling waves across various stimulus configurations. The results align qualitatively with those from a prior fMRI study that investigated traveling waves of binocular rivalry across different attentional states in the human visual cortex (Lee et al., 2007).

Our theoretical model aims to provide mechanistic insights into the generation and propagation of cortical traveling waves and to generate hypotheses regarding their functional relevance in visual perception. Although the phenomenon of perceptual waves has been studied for some time, previous results showing that wave propagation slows down when attention is diverted (Lee et al., 2007) have not yet been accounted for by any existing model. A key novelty of the present model is its ability to capture such attention-related modulations in wave dynamics, thereby offering a theoretical framework to interpret empirical results that have remained partly unexplained. Our model generates experimentally testable predictions and provides a foundation for future studies investigating the role of attention in shaping the dynamics and functional consequences of cortical traveling waves.

## Materials and Methods

### Participants

21 healthy individuals (12 female, 6 left-handed) participated in the experiments with an average age of 24.3 years ± 2.6. Two participants were excluded from the analyses because they did not follow the instructions. All participants had normal or corrected-to-normal vision and reported no history of psychiatric or neurological disorders, gave their written informed consent and were compensated for their participation. All procedures were approved by the French ethics committee Ouest IV - Nantes (IRB #20.04.16.71057) and followed the Code of Ethics of the World Medical Association (Declaration of Helsinki).

### Apparatus and Stimuli

Python (version 3.8.1) and PsychoPy (version 2020.1.2) were used for stimulus display. Stimuli were presented in a dark room at a viewing distance of 110 cm on a gray screen with 1920 × 1080 pixels (210 × 120 cm; 109 × 62 degrees of visual angle, dva) using a PROPixx projector (back-projection) at 120 Hz refresh rate (with DepthQ circular polarizer from VPixx). Blue/Red Filter Glasses were worn by the participants throughout the experiments. The stimuli consisted in two orthogonal gratings (45° and -45° from vertical) windowed by an annulus (gaussian filter applied to inner edge, σ = 0.125 dva) and with different levels of contrast in the two eyes: 80% for the high-contrast eye and 15% for the low-contrast eye). The contrasts were found in pilot experiments to maximize the occurrence of perceptual wave in most of the participants. Annuli could be of one of three different sizes: small (near fovea), medium (near periphery) and large (periphery). The small annulus was defined by an outer radius of 2.8 dva, an inner radius of 0.5 dva and a spatial frequency of 0.77 cycles/degree (cpd); the medium annulus: outer radius = 4.4 dva, inner radius = 1.8 dva, spatial frequency = 0.67 cpd; and the large annulus: outer radius = 6 dva, inner radius = 4.4 dva, spatial frequency = 0.58 cpd. The spatial frequencies were chosen to match cortical magnification (see Broderick et al., 2022).

### Eye-tracking

An EyeLink 1000Plus system ensured participants maintained fixation on a central point. If a fixation break occurred, i.e., if participants’ gaze deviated by >2° or if they blinked, the trial was aborted and the next trial started.

### Psychophysics procedure

The psychophysics experiment replicated and extended two previous studies (Lee et al., 2007; Wilson; 2001). The experiment was split in two sessions of two hours each, one in the morning and one in the afternoon (separate days), to control for potential circadian effects (Zong et al., 2023; Pracucci et al., 2023). The first hour of the first session (morning/afternoon counterbalanced across participants) served as training. Participants then performed three different tasks in separate blocks: (1) main task, (2) diverted attention task, and (3) alternating waves task. At the end of each task, a brief debriefing was conducted to collect participants’ subjective impressions regarding their perceptual experience. One hour of each session was dedicated to the main task, and the remaining hour was split between the two others.

### Main task

The trial sequence started with the presentation of a low-contrast grating annulus to one eye (small, medium or large size; randomly interleaved), followed 30 ms later by a high-contrast grating annulus to the other eye. The delay was intended to facilitate perceptual dominance of the high-contrast annulus (Lee et al., 2007). Participants were instructed to press a key when the high-contrast annulus was dominant. The key press triggered the onset of a brief (100 ms) contrast increment (to 95%) at one of eight equally spaced locations (with an angular range of 30°) of the low-contrast annulus, after a variable delay of 100-250ms (**Figure 1A**). In the event the contrast pulse initiated a perceptual traveling wave, participants were instructed to press a key when the wave reached a target area (marked by dashed lines) at one of the eight equally spaced locations along the annulus (at least 90° between contrast pulse and target area; **Figure 1D**). If the contrast pulse did not evoke a perceptual wave, or if the wave did not reach the target zone, participants did not press any key and the trial automatically ended after 4 seconds. In 20% of the trials (randomly interleaved), the contrast pulse was presented to both eyes at the same location. These double-pulse trials allow us to empirically test whether wave propagation speed changes when both eyes receive an input increment (to 95% of contrast) one of the predictions of our model). Participants performed 24 blocks (12 in the first session and 12 in the second) of 40 trials each (320 trials total per annulus size).

**Figure 1.**
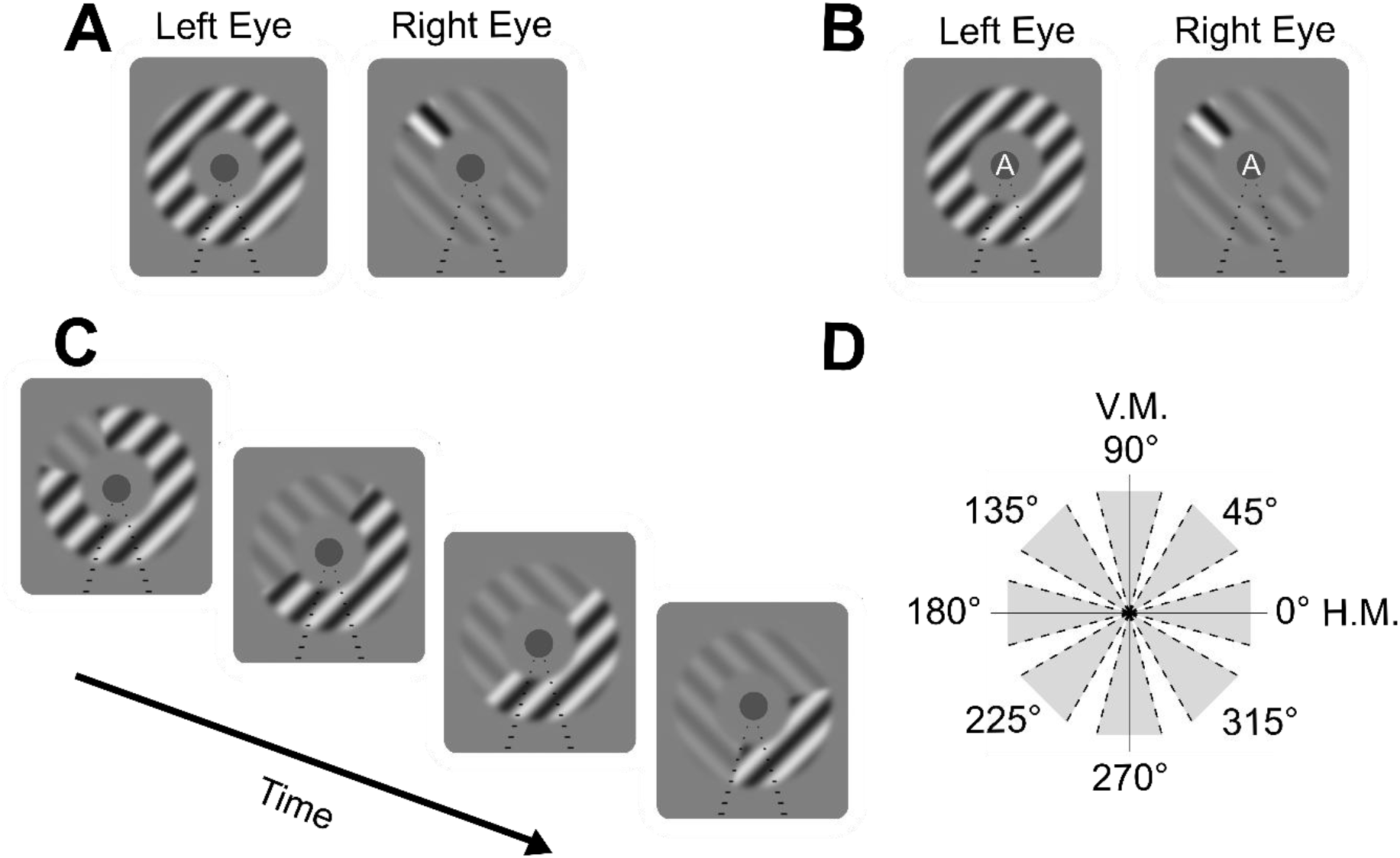
Psychophysics tasks. **(A)** In all tasks, participants were presented with orthogonal grating annuli (one in each eye). A brief 100ms-contrast pulse was presented in the low-contrast grating to trigger a perceptual wave. **(B)** In the diverted attention task, a sequence of white or black letters or numbers (one every 400ms) was presented at fixation after perceptual dominance was established. **(C)** Representation of an illusory perceptual wave. Participants were instructed to press a key when the wave reached the target zone (mark with dashed lines). **(D)** Potential contrast pulse and target zone locations. V.M., vertical meridian; H.M., horizontal meridian.

### Diverted attention task

In the second task, the parameters of the rival gratings and the sequence of events were identical to those in the main task. An additional task was performed at fixation and was aimed at diverting attention from the annulus (**Figure 1B**). A rapid series (one every 400 ms) of small, white or black, letters and/or numbers appeared at fixation after participants pressed a key to indicate stable perceptual dominance. Simultaneous with the letter presentation, the rapid contrast pulse in the low-contrast grating was presented after a variable delay of 100-550 ms (same as in Lee et al., 2007). In addition to the first report (i.e., a perceptual wave reaching the target zone, or no report), participants were instructed to report whether the repeated item in the central stream was a letter or a number. Participants performed 4 blocks of 40 trials each.

### Alternating waves task

This task only used the medium size annulus, and the distance between target area and contrast pulse was kept at 180°. The trial sequence was identical to the main task except that when participants reported the perceptual wave reaching the target area, a second pulse of contrast was presented to the high contrast eye at the target area to induce a perceptual wave traveling in the opposite direction. Participants then pressed a button when this second wave reached the opposite location. Participants performed 4 blocks of 16 trials each, which were used to measure the average propagation time of these alternating perceptual waves. We then used these individually measured propagation times to alternatively present contrast pulses at opposite (180° distance) locations and record how many of these alternating waves could participants perceive until the illusory percept disappeared (participants pressed a key when they did not perceive the illusion any longer). Participants performed 4 blocks of 16 trials each.

### Computational model

Perceptual traveling waves were modeled with a set of equations to simulate the spatio-temporal cortical dynamics of binocular rivalry. The model is intended to characterize neural activity in the visual cortex through signal-processing computations. It is not intended to be a biologically-plausible implementation of the specific underlying circuit, nor of the cellular or molecular processes. Therefore, model elements do not correspond to specific cell types, and parameters such as orientation selectivity, and time constants may emerge from neural circuits rather than being intrinsic neuron features. The model comprises three distinct neural processes/representations: sensory representation, attentional modulation, and mutual inhibition (**Figure 2C**). Each of these processes/representations is repeated in a two-dimensional array of processing modules (**Figure 2A**), with each module corresponding to the hypothesized function of a subpopulation of neurons in a small region of visual cortex. Each module in the array was designed based on the attention model of binocular rivalry from Li et al. (2017). Each module receives an input drive from a specific portion of the visual field (the image) corrected for cortical magnification, with most of the modules receiving input from the central part of the visual field.

**Figure 2.**
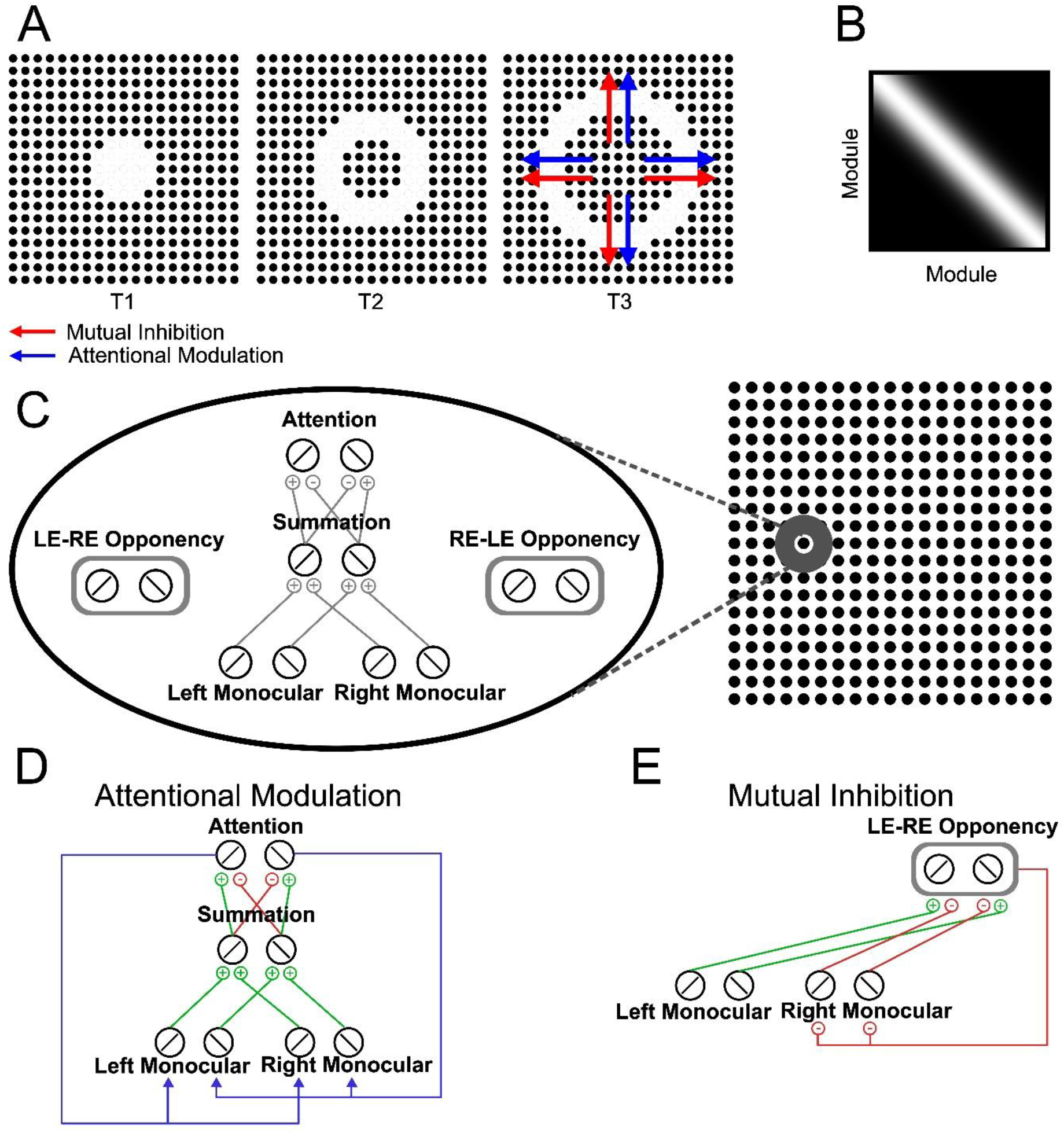
Model of waves in binocular rivalry. **(A)** The model is composed of a series of modules (black dots). Attentional modulation (blue arrows) and mutual inhibition (red arrows) spread across spatially neighboring modules over time (t, time). **(B)** Matrix representing the spatial spread of attention and mutual inhibition (the exponential terms in Equations 2 and 3) as a function of the distance between modules. **(C)** Individual module architecture (from Li et al., 2017). **(D)** Attentional modulation. There are two attention neurons selective for orthogonal orientations. Each attention neuron receives excitatory inputs from the binocular summation neuron that is selective for the same orientation, and suppressive inputs from the orthogonal orientation. Monocular neurons selective for the same orientation receive the same attention gain factor (blue arrows) determined by the response of the attention neuron with the same orientation preference. **(E)** Mutual inhibition. There are two groups of opponency neurons (RE– LE, right-eye – left-eye and LE–RE, left-eye – right-eye). Here, only the RE–LE opponency neurons are illustrated. Opponency neurons compute the response difference between the two eyes for a particular orientation. The LE–RE (RE–LE) monocular neurons are inhibited by the RE–LE (LE–RE) opponency neurons. For illustration purposes, in (D) and (E) the feedback from the attention neurons or opponency neurons are only illustrated within the local module. In the full model, feedback spread across modules as in (A).

In terms of sensory representation, each possible stimulus location in the model includes six neurons: two neurons for left-eye monocular processing, two for right-eye monocular processing, and two that performed binocular summation. Each pair of neurons are selective to orthogonal orientations.

The response of the left-eye monocular neuron R_l1i_ is computed as:

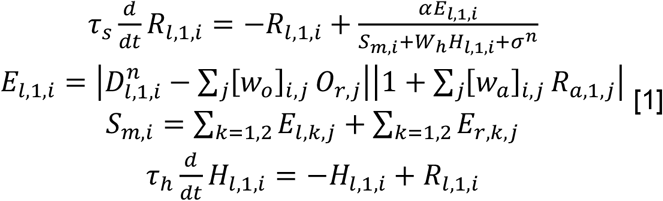

Here l and r specify left-eye and right-eye respectively, subscripts 1 and 2 specify the two orientations, and j and i denote locations. The first line is an equation for calculating the response over time in terms of excitatory drive (E), suppressive drive (S), and adaptation (H). The subsequent lines provide expressions for E, S and H. The excitatory drive (E) is determined by the input (D) that is modulated by two factors. First, the pooled responses of the opponency neurons (O_r_) that responded to the opposite eye are subtracted from the input D. The strength of this subtractive suppression is determined by [W_o_]_ij_ representing the weight of suppression from the j^th^ opponency population to the i^th^ monocular population. Second, the resulting activity after the subtractive suppression is multiplied by an attention gain factor |1 + ∑_*j*_[*w*_*a*_]_*i,j*_ *R*_*a*,1,*j*_|, in which 1 is the baseline of attention gain, and R_a1j_ is the response of the attention neuron that is selective for the same orientation at location j. The strength of attentional modulation is determined by [W_a_]_ij_ representing the weight of attentional feedback from the j^th^ attention population to the i^th^ monocular population.

The interactions between neighboring (spatially adjacent) modules are mediated through the spatial spread of mutual inhibition (W_o_) and attentional modulation (W_a_). Each module is located at a 2D position r_i_ = (x_i_, y_i_) on a simulated cortical surface. We define the Euclidean distance between two modules as *d*_ij_ = ‖*r*_i_ − *r*_j_‖ = √(*x*_i_ − *x*_j_)^2^ + (*y*_i_ − *y*_j_)^2^. The weights [W_o_]_ij_ and [W_a_]_ij_ decay exponentially with distance (**Figure 2B**). The weight of mutual inhibition is:

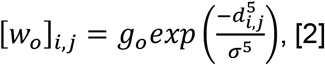

in which g_o_ controls the strength of mutual inhibition. The weight of attentional modulation is

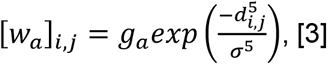

in which g_a_ controls the strength of attention. We use a single parameter σ to determine the spatial extent of mutual inhibition and attention. The choice of the exponent 5 plays a critical role in shaping the effective spatial influence of attention and inhibition and was motivated empirically to balance the extent of the effect across modules. Lower exponents led to overly broad interactions and abolished wave dynamics, while higher exponents restricted the spread too narrowly to allow propagation. An exponent of 5 thus allowed stable wave behavior with biologically plausible ranges for the other parameters, consistent with similar formulations used in prior models of perceptual wave propagation (Wilson et al., 2001).

Except for those above, model components (binocular-summation, opponency and attention neurons) are implemented in the same way as in Li et al. (2017).

Model variants were developed to assess different dynamics observed in other binocular rivalry experiments, thus exploring the robustness of the theoretical framework. These include adjustments for traveling wave’s speeds in response to collinear patterns, implemented by adding a collinear facilitation term as described in Wilson et al. (2001) and Kang et al. (2010):

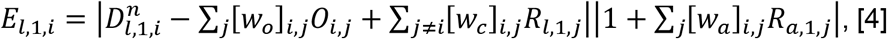

in which:

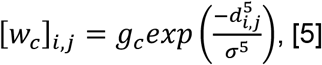

where *w*_*c*_ determines the spatial spread and the strength of collinear facilitation. The value of g_c_ is set to zero when simulating the traveling wave for a non-collinear (radial grating) stimulus, and to 0.32 when simulating the traveling wave for a collinear (concentric grating) stimulus. To simulate effects of recurrent excitation, which enhances the excitatory drive of monocular neurons, we adopted a similar formulation:

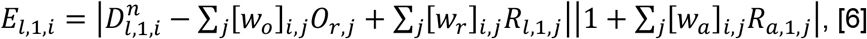

in which

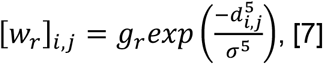

where *w*_*r*_ determines the strength of recurrent excitation. Here, the recurrent excitation includes the response of the neuron itself while the collinear facilitation is driven by responses of other neurons. Recurrent excitation has the same strength in both eyes whereas collinear facilitation enhances activity only for the neural representation corresponding to the eye presented with the collinear pattern.

In another model variant, we simulated recurrent excitation that was purely local. In this case, each neuron received recurrent input only from itself, and the excitatory drive of the monocular neurons was computed as:

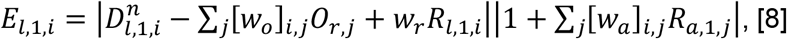

The values of the parameters are based on the previous study from Li et al. (2017) with some modifications. Modified parameters are reported in **Table 1**.

**Table 1.**
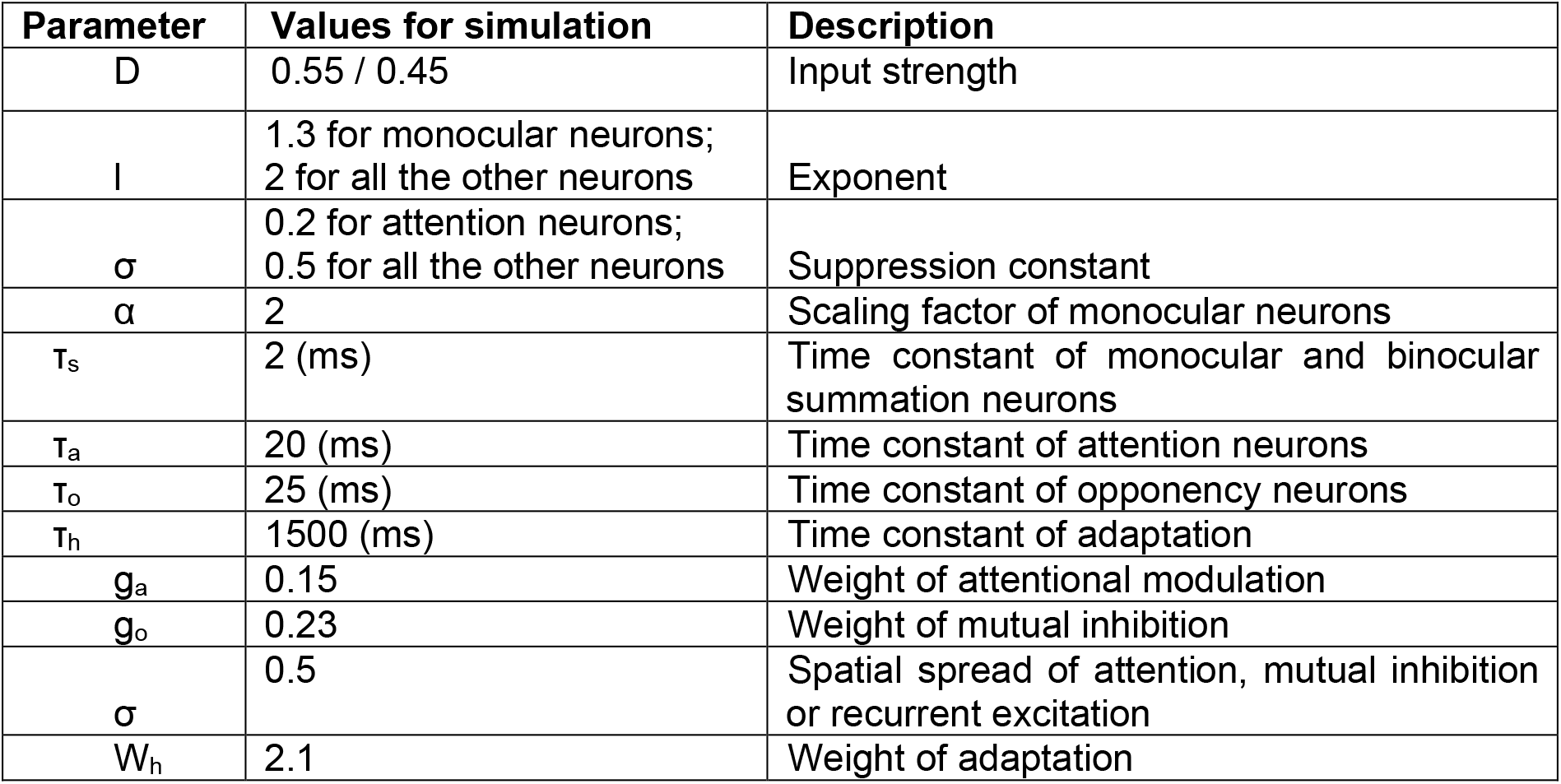
Model parameters.

To quantify speed, the model required assumptions about the physical spacing in cortex and number of modules. We assumed that the distance between spatially adjacent modules was 2.5 mm (based on Wandell & Winawer 2015 and measures of population receptive fields, pRFs). To simulate a participant’s V1 activity, the model generated a random polygon with an adjustable area size. The polygon was then rescaled to match a realistic area size of a human V1 cortex (here 2000 mm2; Benson et al., 2022) and further segmented in modules (separated by 2.5 mm) corrected for cortical magnification.

## Data Availability

The dataset generated in the current study, as well as experiment, analysis and modeling codes will be made available upon publication.

## Results

### Psychophysics

Participants performed a series of tasks under binocular rivalry, in which a brief contrast increment was used to induce a change in dominance that started at one location in the visual field and expanded progressively, like a wave, as the dominant annulus became suppressed. We measured the traveling properties of these perceptual waves across different manipulations, including different stimulus sizes and angular distances between the location of the contrast increment and the location of a target zone (**Figure 1A**, participants pressed a key when the perceptual wave reached the target, dashed area). This task provided an ideal tool for acquiring data to fit our model and to test key properties of wave propagation, such as asymmetries across the visual field, interhemispheric transfer, and the ability to exclude reaction times not related to wave dynamics from our measurements.

We first assessed for all combinations of pairs of angular distances whether response times varied as a function of visual field locations (**Figure 1D**) and annulus sizes, but there was no significant effect (Kruskal-Wallis test, n.s.). This result suggests that wave propagation was not detectably influenced by quadrant or hemispheric location in our experiment.

Given this absence of spatial asymmetries, we report the response times aggregated for the three possible distances, i.e., 90°, 135° and 180° regardless of position, separately for the three stimulus sizes, i.e., small, medium and large (**Figure 3**). Response times increased with both distance to travel (Kruskal-Wallis test for distance of 90°: H-statistic: 28.5, p-value: <0.001; 135°: H-statistic: 73.7, p-value: <0.001; 180°: H-statistic: 15.6, p-value: <0.001) and annulus size (small annulus: H-statistic: 100.1, p-value: <0.001; medium: H-statistic: 115.8, p-value: <0.001; large: H-statistic: 133.7, p-value: <0.001). The response time increase, however, was not directly proportional to the travel distance. This may be due to delays in motor responses and/or decision biases affected by the distance to travel. Consequently, we then considered the response time differences between the smallest (90°) and largest (180°) distance, thus allowing for a less biased measure. The differences of response times displayed a remarkable consistency across conditions: 316 ms for the small annulus, 325 ms for the medium, and 334 ms for the large. Angular speeds, for each annulus, were: 284°/s for the small annulus, 276°/s for the medium, and 269°/s for the large.

**Figure 3.**
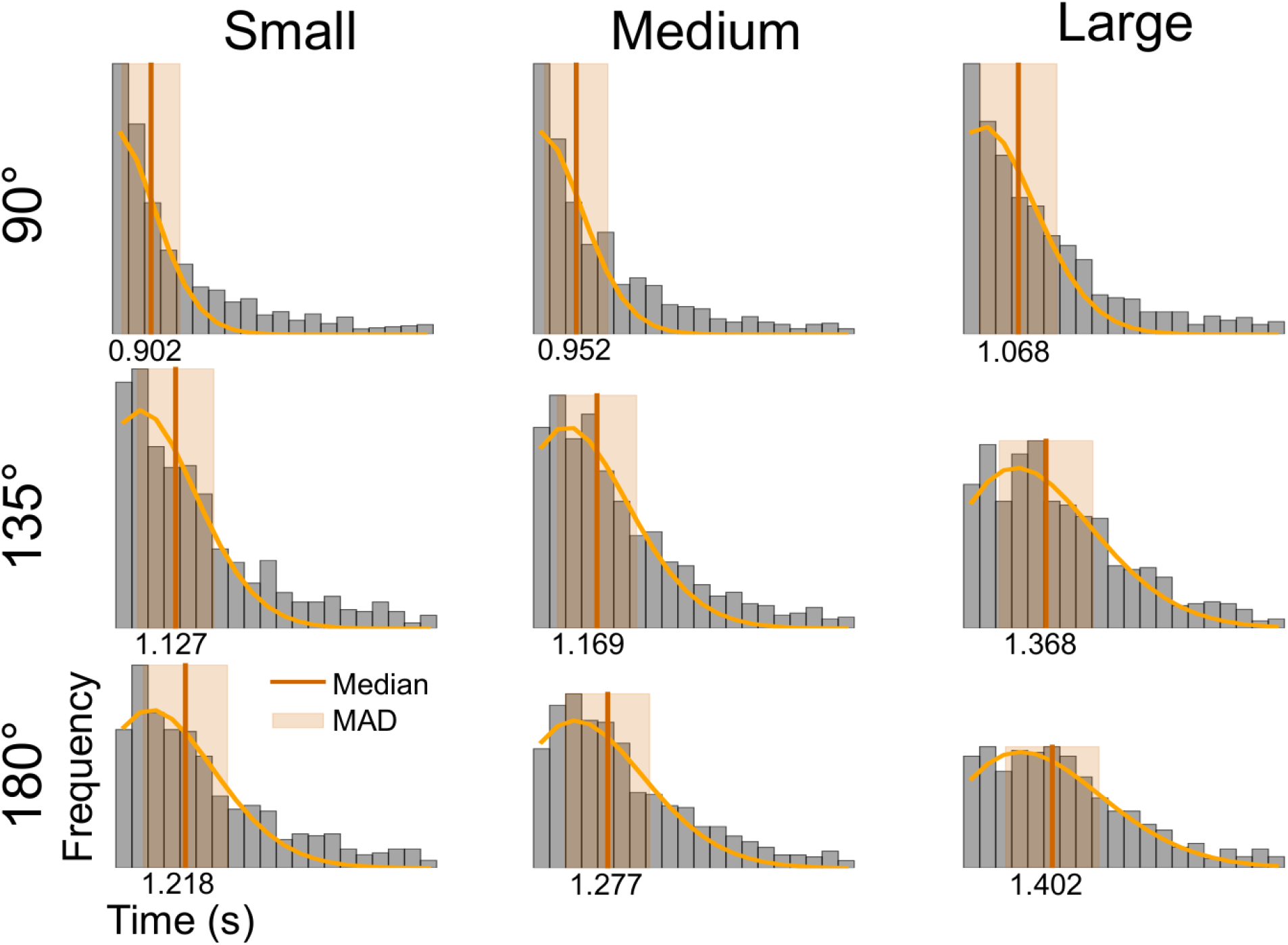
Behavioral results. Histograms show the distribution of response times across participants, with medians and median absolute deviations (MAD) for each condition (annulus size: small, medium and large; distances until target zone: 90°, 135° and 180°).

Note that there was a small but significant difference in reaction times between participants performing the experiment in the morning versus in the afternoon (Mann-Whitney U test statistic: 3229825, p-value: <0.001; 1.11s on average for morning; 1.28s for afternoon) suggesting a circadian modulation of perceptual wave propagation.

On average, there was no significant difference across stimulus size in the number of trials for which a perceptual wave was not perceived (Kruskal-Wallis H-statistic: 0.866, p-value: 0.648), although there was substantial variability across individual participants (**Figure 4**).

**Figure 4.**
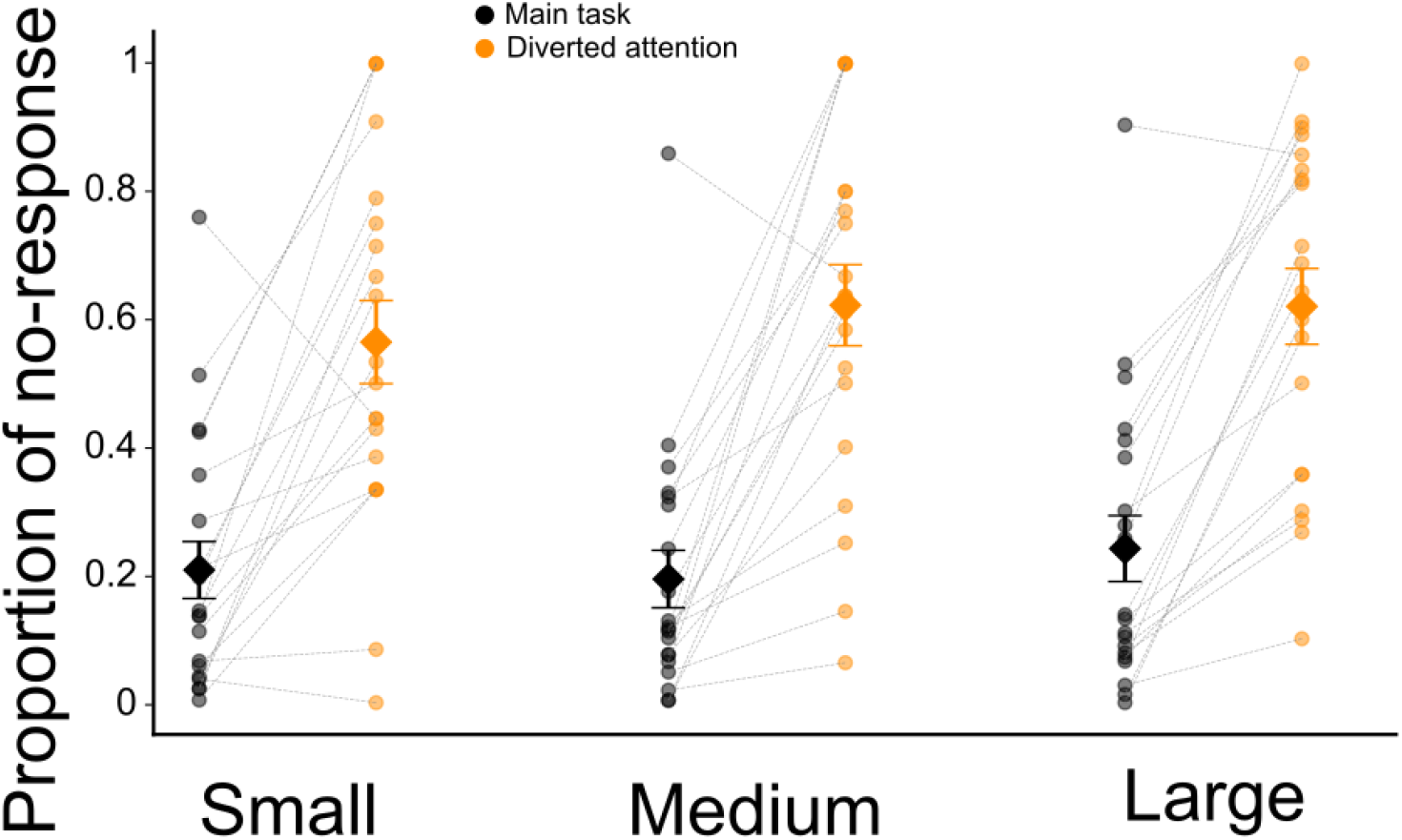
Proportion of trials with no perceived waves, with or without attention. Lights dots, individual participants. Black diamonds, average across participants. Error bars: Standard Error of the Mean (SEM).

In the *diverted attention* task, participants’ attention was maintained at the center of the screen, away from the peripheral annulus. Participants were instructed to report whether the repeated item in the central stream (**Figure 1B**) was a letter or a number. Diverting participants’ attention from the principal task (pressing a button when the perceptual wave reaches target zone) offered an ideal way to assess the role of attention in this perceptual illusion. The resulting behavioral changes strongly support the model’s prediction of a high sensitivity to attentional modulation in wave dynamics. Average performance was 66% ± 8% of correct responses (significantly above chance, Wilcoxon signed-rank test statistic: 13.0, p-value: <0.001) suggesting that participants successfully maintained attention at fixation. Importantly, in this task, participants reported significantly fewer perceptual waves, i.e., mean: 39.5%, SEM: 5.9%; versus mean: 78.6%, SEM: 4.6% in the previous task (Wilcoxon signed-rank test statistic: 6, p-value: <0.001; **Figure 4**). These results are consistent with previous findings reporting that attention is necessary for perceptual traveling waves to emerge (Lee et al., 2007).

Post-task debriefings confirmed that participants perceived fewer waves and when they did perceive a change, they often described the perceptual transitions as abrupt or incomplete. They also reported lower confidence in judging whether the wave had reached the target zone. These subjective impressions align with the reduced wave reports (**Figure 4**), suggesting a shift from gradual wave propagation to more instantaneous perceptual switches under diverted attention.

Finally, in the *alternating waves task*, participants were presented with flash increments presented periodically at opposite locations in each eye alternatively (delays between flashes adjusted for each individual; see Methods). This task allowed us to test a new experimental condition predicted by our model, and not previously reported in the literature, showing the robustness of our theoretical framework. On average, participants perceived 4±2 waves before the percept faded away, showing the consistency of the delays of wave propagation over time.

### Model

We simulated neural responses corresponding to the measured perceptual traveling wave speeds.

**Movie 1.**
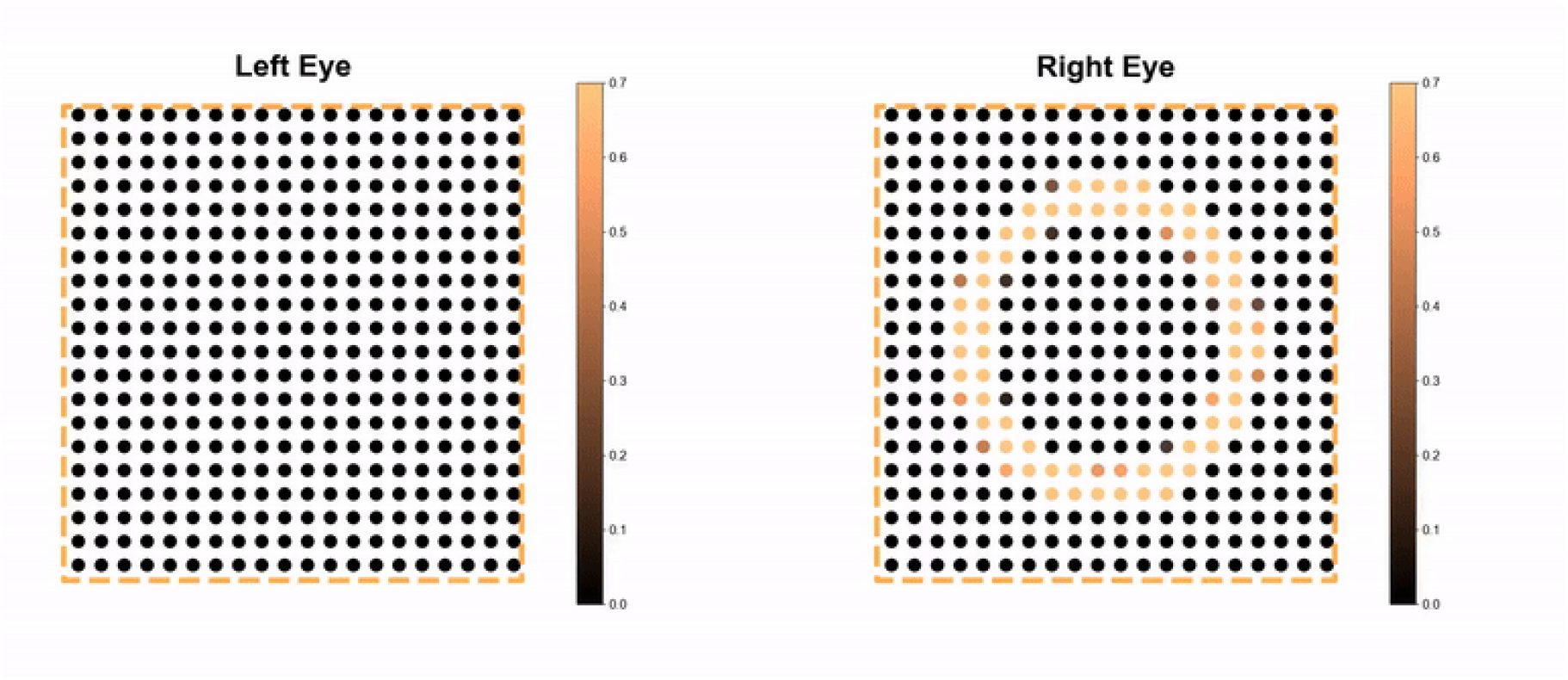
Module activation in response to stimulus.

The model exhibited wave propagation in response to the contrast (trigger) increment. The suppressed stimulus broke through the suppression at the trigger location and propagated across the retinotopic representation of the annulus (**Movie 1**). The responses of neurons that prefer the orientation of the dominant stimulus were initially high and exhibited a progressive decrease in activity after the contrast increment, while inversely, the responses of neurons that prefer the other orientation were initially low and exhibited a progressive increase in activity.

We varied the strength of attentional modulation (τ_a_) and simulated traveling waves under various levels of attention. We found that the wave’s speed decreased as the strength of attentional modulation increased (**Figure 5**). In the model, the effect of attention on wave speed was similar to that of mutual inhibition (mediated through opponency neurons), i.e., increasing the strength of mutual inhibition (*τ*_*o*_) slowed down the propagation.

**Figure 5.**
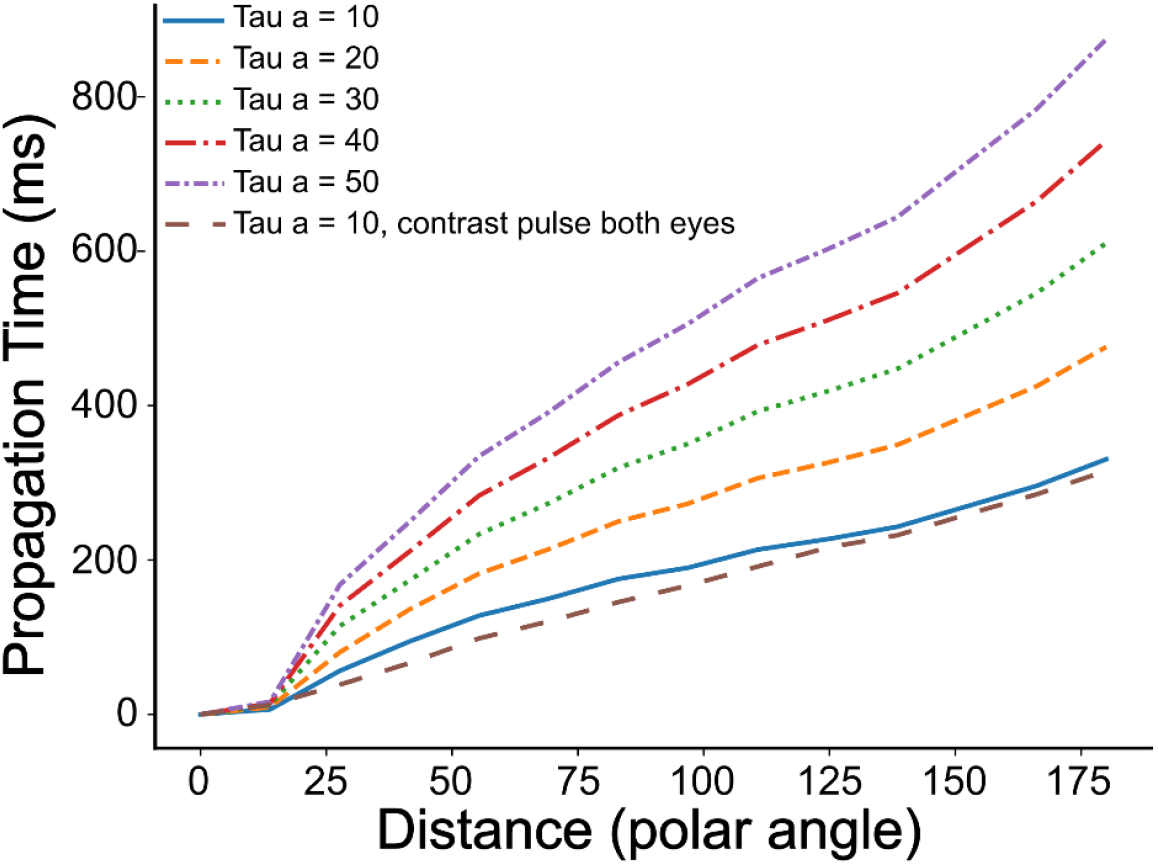
Effect of attentional gains on simulated traveling waves. Cumulative distance (in polar angle) traveled by the wave as a function of time. Each line in the plot is an individual simulation with a varying attentional modulation (τ_a_). Dash brown line in the plot: simulation where a contrast pulse was presented to both eyes, simultaneously.

To extract the propagation time from the simulated neural responses, we then read out the first time point (after the trigger) at which the response to the low contrast annulus exceeded the response of the high contrast one, for each module separately. When reporting propagation speed, we only considered the time point at which the perceptual wave reached the module corresponding to the retinotopic location of the annulus target zone. The model can fit the behavioral measures of perceptual wave’s speed. An example of a simulation can be seen in **Figure 6**.

**Figure 6.**
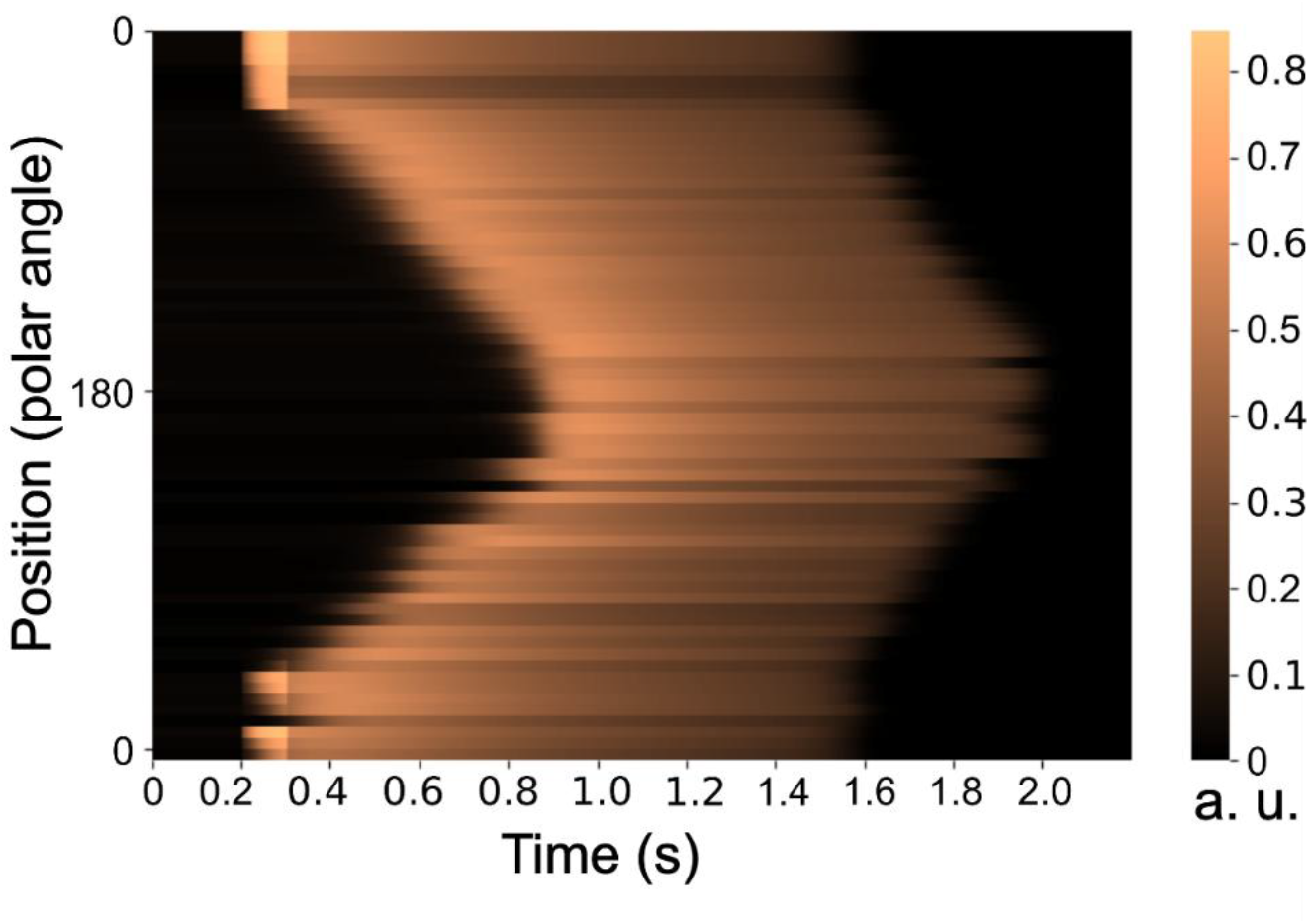
Simulation of traveling wave propagation for the large annulus. Activity for all activated modules (y-axis) organized by polar angle position relative to the annulus. Contrast pulse increment is observed from 200 to 300ms and generate a wave across the simulated annulus reproducing the perceptual wave’s speed observed in psychophysics of 334ms for a 90° distance.

Traveling wave speed has been reported to be faster when the stimulus has a collinear structure (Alais & Blake, 1999; Kang & Blake, 2011; Kang et al., 2010; Wilson et al., 2001). Our model accounted for this speed increase by adding collinear facilitation between neighboring sensory neurons (**Figure 7A**), consistent with previous models (Kang et al., 2010; Wilson et al., 2001).

**Figure 7.**
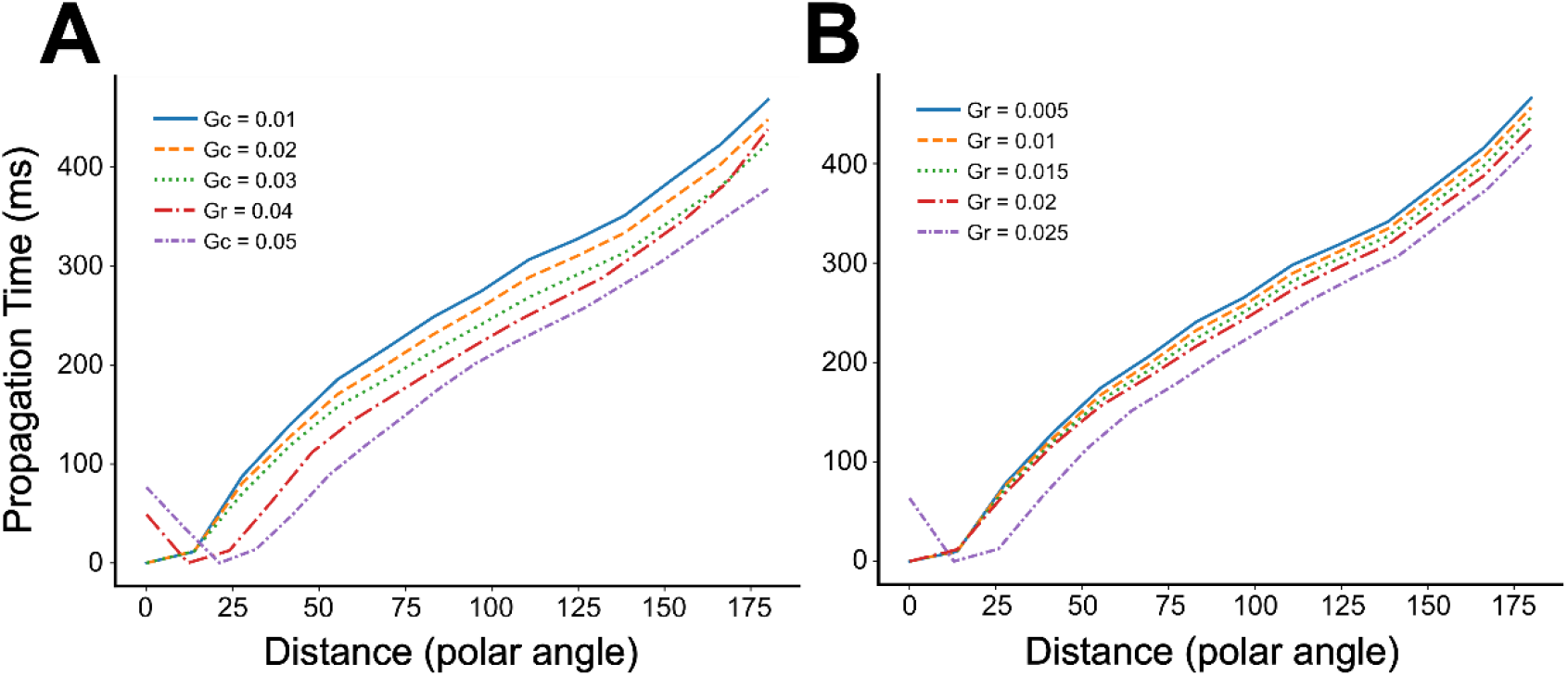
Effect of collinear facilitation and recurrent excitation on simulated traveling waves. **(A)** The effect of collinear facilitation on wave speed for a constant attention level (τ_a_=20ms). Each line corresponds to a different strength of collinear facilitation (g_c_ see Equation 4). **(B)** Same as (**A**) except that each line corresponds to a different strength of recurrent excitation (g_r_, see Equation 6) coming from neighboring modules.

Recurrent excitation also altered wave speed, depending on its spatial extent. When recurrent excitation was included in the model, across spatially adjacent modules (Equation 3; **Figure 7B**), it accelerated the propagation, consistent with an earlier study (Bressloff & Webber, 2014). When recurrent excitation was only local (Equation 8), the speed did not change. When the recurrent excitation was high, the model stayed in a pure winner-take-all regime as the high-contrast stimulus maintained dominance with no response alternation even in the presence of the trigger.

In our diverted attention experimental protocol, we found that traveling waves were seen less frequently (39% of trials) and, when seen, much faster than in the main task (median value of 0.22 s, against 1.17 s). Also, previous studies done by Lee et al. (2005, 2007) found that the peak of the fMRI responses reflected a traveling wave of neuronal activity and the traveling wave speed, inferred from the fMRI response latencies, was slower in the focused-attention condition than in the diverted-attention condition. Model simulations yielded analogous results; traveling waves were slower when the attention gain was smaller, as shown in **Figure 5**.

Finally, our model could reproduce the alternating waves without further parameter change (**Movie 2**). However, the model did not capture appropriately the output of the condition in which a contrast increment was applied to both eyes simultaneously (modeled by increasing the input strength D for both eyes). Although the behavioral results showed a decrease in propagation speed (130ms decrease on average), the model instead predicted a speed increase (23ms faster).

**Movie 2.**
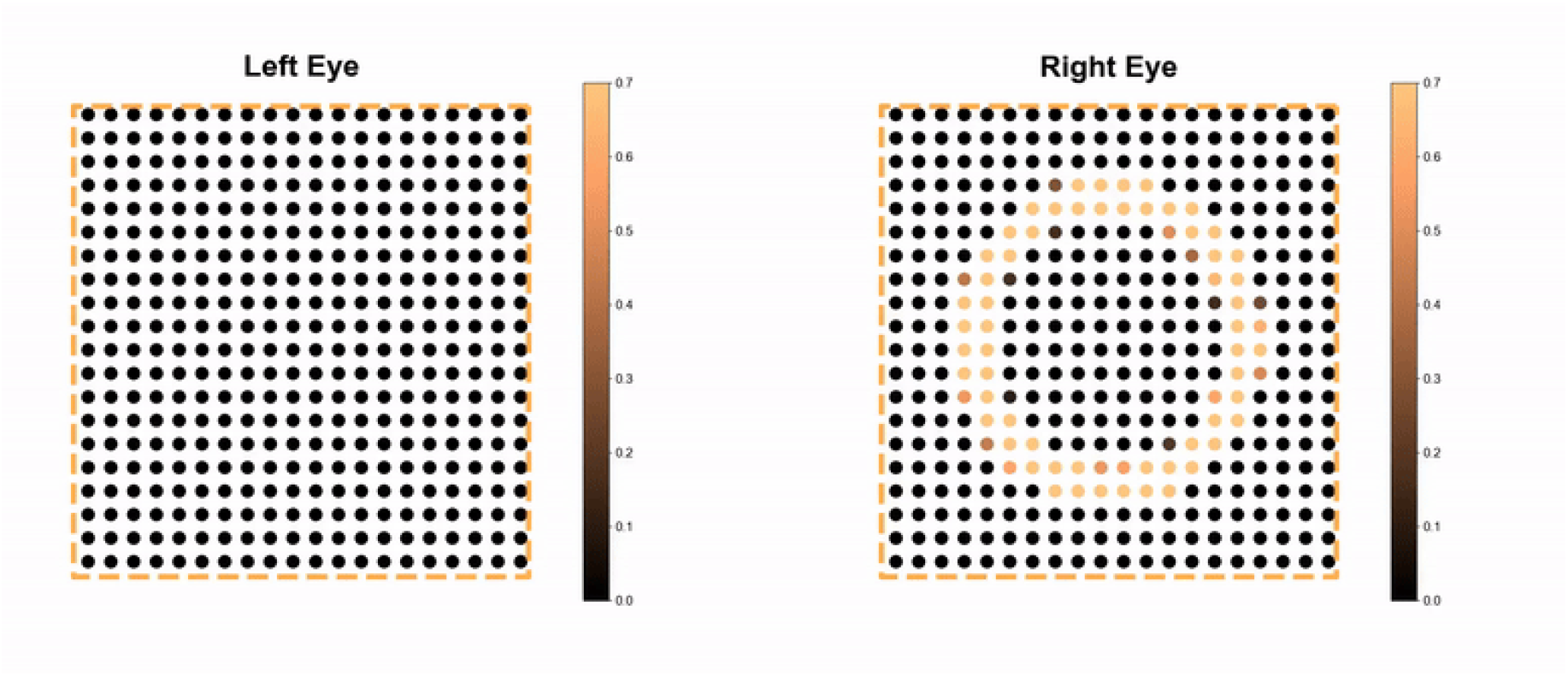
Periodic Traveling waves.

## Discussion

Studying cortical waves traveling across short, mesoscopic distances in the healthy human brain is constrained by several methodological considerations, especially the poor spatial resolution of available non-invasive recording techniques (Grabot et al., 2024; Petras et al., 2025). We employed a dual approach of psychophysics and computational modeling to address this issue. We used the illusory phenomenon of perceptual waves in binocular rivalry (Wilson et al., 2001) to investigate the mechanisms underlying wave propagation in retinotopic areas. Replicating previous psychophysics experiments, perceptual waves propagate across the retinotopic space with durations increasing with distance and eccentricity (Wilson et al., 2001). Diverted attention significantly reduced the occurrence and duration of the waves, replicating previous work suggesting that attention plays a pivotal role in the phenomenon (Lee et al., 2007). Our computational model successfully replicated the key dynamics of perceptual waves, demonstrating the role of attentional modulation and mutual inhibition in wave propagation, as well as the influence of collinear facilitation and recurrent excitation. The model also accounted for variations in wave propagation for different distances and eccentricities, commensurate with cortical magnification. Together, our results bridge the gap between observed perceptual phenomena and underlying neural mechanisms, offering insights into visual cortical processing and guiding future experimental investigations.

Our experimental design allowed us to test several additional questions. First, we assessed potential variations in propagation times due to visual field asymmetries or interhemispheric crossing. A vertical meridian crossing penalty, in terms of propagation time, was expected due to interhemispheric communication, but our results did not reveal significant differences between intra- and inter-hemispheric propagation. Previous studies have reported mixed results on this question: while some authors found no significant effect (Genc et al., 2011), others did report a measurable delay for interhemispheric propagation (Wilson et al., 2001). Additionally, during debriefing, several participants reported that waves were easier to perceive in the lower visual field, in line with known visual field asymmetries in perceptual performance (e.g., Barbot et al., 2021). These subjective reports suggest that field-dependent differences may influence perceptual propagation, although they were not captured in our behavioral measures. Further experimentation is thus warranted.

It is important to highlight that our empirical results relied on a measure of reaction times to assess the speed of the perceptual wave. Reaction times, however, are known to be compound measures of sensitivity and decision criteria and include motor response times (Kienitz et al., 2021). They are thus less sensitive to small changes between conditions, which would explain the lack of reaction time difference, especially within versus between hemifield propagation. This is not an issue for the main outcome of the experiments as we compared reaction times between conditions with different propagation distances and stimulus eccentricities. The results of this manipulation suggested that the spatial scaling of neural representations in V1 helps maintain temporal consistency across varying eccentricities.

The results further show substantial inter-individual variability in wave perception. For some participants, the percentage of perceived waves for a given eccentricity was as low as 10% of trials. It appeared in the post-experiment debriefing that difficulties in perceiving waves under certain conditions could arise from challenges in maintaining the high-contrast annulus in a stable perceptual state long enough to experience the illusion. Interestingly, our model effectively reproduced the empirical propagation times. It can be applied to fit either a general distribution of propagation times or participant-specific values. Further investigation will focus on using the model to further assess inter-individual variabilities in this perceptual wave phenomenon. Inter-individual variability was additionally observed between participants who performed the experiment in the morning versus those who participated in the afternoon. There are two alternative explanations for such a result. This could either be due to modulation in vigilance or in arousal due to circadian rhythms, both modulating attention necessary for the perceptual wave to emerge. Further experimentation is required to disentangle the two interpretations. The proposed model effectively reproduced the observed propagation times and offers the potential to predict as-yet-unobserved behaviors.

Some limitations, however, warrant further investigation. First, the model in its current form fails to exhibit wave propagation when the inputs for the low-contrast and high-contrast stimulus differ by more than a small margin, a constraint that does not align with the experimental manipulation that was necessary to stabilize the first dominant percept. Second, the model predicts that increasing the input strength for both eyes accelerate the speed of the wave, which was not supported by our results. These discrepant results can be explained by the simplified manner in which the model currently handles input, relying solely on arrays of contrast values. Future work will address these concerns by integrating more biologically grounded mechanisms into the model. Specifically, we will build on the proposal of a recurrent circuit model between LGN cells, principal cells and modulator cells in V1, which was shown to produce normalization and replicate known properties of V1 dynamics (Heeger & Zemlianova, 2020).

## Conclusion

Our study proposes a model of attention-induced, perceptual traveling waves across the retinotopic space. While previous work relied on external stimuli to induce propagating cortical activity (Sokoliuk and VanRullen, 2016; Fakche & Dugué, 2024), we use a paradigm in which the perceptual wave is illusory and develop a computational model to provide new insights into the underlying mechanisms.

## Acknowledgements

This project has received funding from the European Research Council (ERC) under the European Union’s Horizon 2020 research and innovation programme (grant agreement No 852139 - Laura Dugué).

## References

Alais, D., & Blake, R. (1999). Grouping visual features during binocular rivalry. Vision Research, 39(25), 4341–4353. 10.1016/S0042-6989(99)00146-7

Alamia, A., & VanRullen, R. (2019). Alpha oscillations and traveling waves: Signatures of predictive coding? PLOS Biology, 17(10), e3000487. 10.1371/journal.pbio.3000487

Alexander, D. M., & Dugué, L. (2024). The dominance of global phase dynamics in human cortex, from delta to gamma. eLife, 13, RP100674. 10.7554/eLife.100674.1

Alexander, D. M., Jurica, P., Trengove, C., Nikolaev, A. R., Gepshtein, S., & van Leeuwen, C. (2013). Traveling waves and trial averaging: The nature of single-trial and averaged brain responses in large-scale cortical signals. NeuroImage, 73, 95–112. 10.1016/j.neuroimage.2013.01.016

Barbot, A., Xue, S., & Carrasco, M. (2021). Asymmetries in visual acuity around the visual field. Journal of Vision, 21(1), 2. 10.1167/jov.21.1.2

Benson, N. C., Jamison, K. W., Arcaro, M. J., et al. (2022). Variability of the surface area of the V1, V2, and V3 maps in a large sample of human observers. Journal of Neuroscience, 42(43), 8629–8646. 10.1523/JNEUROSCI.0690-21.2022

Blake, R., & O’Shea, R. P. (2009). Binocular rivalry. In L. R. Squire (Ed.), Encyclopedia of Neuroscience (Vol. 2, pp. 179–187). Academic Press.

Blake, R. (2022). The perceptual magic of binocular rivalry. Current Directions in Psychological Science, 31(2), 126–133. 10.1177/09637214211057564

Brascamp, J. W., & Blake, R. (2012). Inattention abolishes binocular rivalry: Perceptual evidence. Psychological Science, 23(10), 1159–1167. 10.1177/0956797612440100

Brascamp, J. W., van Ee, R., Pestman, W. R., & van den Berg, A. V. (2005). Distributions of alternation rates in various forms of bistable perception. Journal of Vision, 5(4), 1. 10.1167/5.4.1

Bressloff, P. C., & Webber, M. A. (2012). Neural field model of binocular rivalry waves. Journal of Computational Neuroscience, 32(2), 233–252. 10.1007/s10827-011-0351-y

Broderick, W. F., Simoncelli, E. P., & Winawer, J. (2022). Mapping spatial frequency preferences across human primary visual cortex. Journal of Vision, 22(4), 3. 10.1167/jov.22.4.3

Fakche, C., & Dugué, L. (2024). Perceptual cycles travel across retinotopic space. Journal of Cognitive Neuroscience, 36(2), 200–216. 10.1162/jocn_a_02075

Genc, E., Bergmann, J., Tong, F., Blake, R., Singer, W., & Kohler, A. (2011). Callosal connections of primary visual cortex predict the spatial spreading of binocular rivalry across the visual hemifields. Frontiers in Human Neuroscience, 5, 161. 10.3389/fnhum.2011.00161

Grabot, L., Merholz, G., Winawer, J., Heeger, D. J., & Dugué, L. (2025). Traveling waves in the human visual cortex: An MEG-EEG model-based approach. PLOS Computational Biology, 21(4)2.e1013007. 10.1371/journal.pcbi.1013007

Hangya, B., Borhegyi, Z., Szilágyi, N., Freund, T. F., & Varga, V. (2009). GABAergic neurons of the medial septum lead the hippocampal network during theta activity. Journal of Neuroscience, 29(25), 8094–8102. 10.1523/JNEUROSCI.5665-08.2009

Heeger, D. J., & Zemlianova, K. O. (2020). A recurrent circuit implements normalization, simulating the dynamics of V1 activity. Proceedings of the National Academy of Sciences, 117(37), 22494–22505. 10.1073/pnas.2004299117

Hindriks, R., van Putten, M. J. A. M., & Deco, G. (2014). Intra-cortical propagation of EEG alpha oscillations. NeuroImage, 103, 444–453. 10.1016/j.neuroimage.2014.08.005

Hughes, J. R. (1995). The phenomenon of travelling waves: A review. Clinical Electroencephalography, 26(1), 1–6. 10.1177/155005949502600101

Kang, M.-S., & Blake, R. (2011). An integrated framework of spatiotemporal dynamics of binocular rivalry. Frontiers in Human Neuroscience, 5, 88. 10.3389/fnhum.2011.00088

Kang, M.-S., Lee, S.-H., Kim, J., Heeger, D. J., & Blake, R. (2010). Modulation of spatiotemporal dynamics of binocular rivalry by collinear facilitation and pattern-dependent adaptation. Journal of Vision, 10(5), 3. 10.1167/10.5.3

Lee, S.-H., Blake, R., & Heeger, D. J. (2003). Traveling waves of activity in V1 correlate with perceptual dominance during binocular rivalry. Journal of Vision, 3(9), 8. 10.1167/3.9.8

Lee, S.-H., Blake, R., & Heeger, D. J. (2007). Hierarchy of cortical responses underlying binocular rivalry. Nature Neuroscience, 10, 1048–1054. 10.1038/nn1939

Lee, S.-H., Blake, R., & Heeger, D. J. (2005). Traveling waves of activity in primary visual cortex during binocular rivalry. Nature Neuroscience, 8, 22–23. 10.1038/nn1365

Levelt, W. J. (1965). On binocular rivalry. Institute for Perception RVO-TNO.

Li, H.-H., Rankin, J., Rinzel, J., Carrasco, M., & Heeger, D. J. (2017). Attention model of binocular rivalry. Proceedings of the National Academy of Sciences, 114(25), E6192–E6201. 10.1073/pnas.1620475114

Logothetis, N. K. (1998). Single units and conscious vision. Philosophical Transactions of the Royal Society B: Biological Sciences, 353, 1801–1818. 10.1098/rstb.1998.0333

Muller, L., Piantoni, G., Koller, D., et al. (2016). Rotating waves during human sleep spindles organize global patterns of activity that repeat precisely through the night. eLife, 5, e17267. 10.7554/eLife.17267

Muller, L., Chavane, F., Reynolds, J., & Sejnowski, T. J. (2018). Cortical travelling waves: Mechanisms and computational principles. Nature Reviews Neuroscience, 19, 255–268. 10.1038/nrn.2018.20

Pearson, J., & Brascamp, J. (2008). Sensory memory for ambiguous vision. Trends in Cognitive Sciences, 12(9), 334–341. 10.1016/j.tics.2008.05.006

Petras, K., Grabot, L., & Dugué, L. (2025). Locally induced traveling waves generate globally observable traveling waves. bioRxiv. 10.1101/2025.01.07.630662

Kienitz, R., Schmid, M. C., & Dugué, L. (2022). Rhythmic sampling revisited: Experimental paradigms and neural mechanisms. European Journal of Neuroscience, 55(11–12), 3010– 3024. 10.1111/ejn.15489

Pracucci, E., Graham, R. T., Alberio, L., et al. (2023). Daily rhythm in cortical chloride homeostasis underpins functional changes in visual cortex excitability. Nature Communications, 14, 7108. 10.1038/s41467-023-42711-7

Sokoliuk, R., & VanRullen, R. (2016). Global and local oscillatory entrainment of visual behavior across retinotopic space. Scientific Reports, 6, 25132. 10.1038/srep25132

VanRullen, R. (2016). Perceptual cycles. Trends in Cognitive Sciences, 20(10), 723–735. 10.1016/j.tics.2016.07.006

Wandell, B. A., & Winawer, J. (2015). Computational neuroimaging and population receptive fields. Trends in Cognitive Sciences, 19(6), 349–357. 10.1016/j.tics.2015.03.009

Wilson, H. R., Blake, R., & Lee, S.-H. (2001). Dynamics of travelling waves in visual perception. Nature, 412, 907–910. 10.1038/35091066

Zhang, P., Jamison, K., Engel, S., He, B., & He, S. (2011). Binocular rivalry requires visual attention. Neuron, 71(2), 362–369. 10.1016/j.neuron.2011.05.035

Zong, F. J., Min, X., Zhang, Y., Li, Y. K., Zhang, X. T., Liu, Y., & He, K. W. (2023). Circadian time- and sleep-dependent modulation of cortical parvalbumin-positive inhibitory neurons. EMBO Journal, 42, e111304. 10.15252/embj.2022111304

