## Supplementary figures and images for "Attention induced perceptual traveling waves in binocular rivalry"

### Movie 1

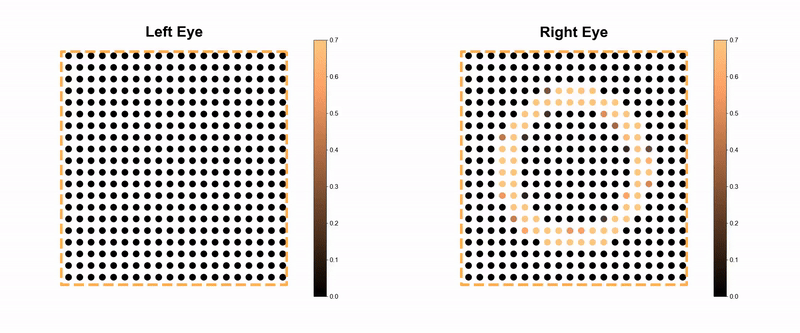

### Movie 2

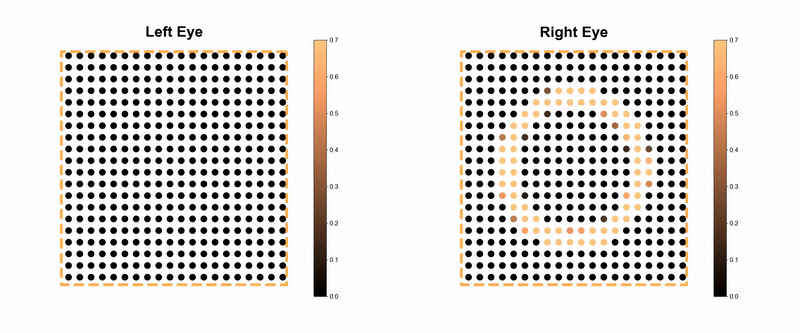
